# Brain-wide imaging of an adult vertebrate with image transfer oblique plane microscopy

**DOI:** 10.1101/2022.05.16.492103

**Authors:** Maximilian Hoffmann, Jörg Henninger, Lars Richter, Benjamin Judkewitz

**Author notes:** These authors contributed equally to this work (Maximilian Hoffmann, Jörg Henninger).

## Abstract

Optical imaging is a powerful tool to visualise and measure neuronal activity. However, due to the size and opacity of vertebrate brains it has until now been impossible to simultaneously image neuronal circuits at cellular resolution across the entire adult brain. This is true even for the smallest known vertebrate brain in the teleost *Danionella*, which is still too large for existing volumetric imaging approaches. Here we introduce image transfer oblique plane microscopy, which uses a new optical refocusing solution via a custom fibre-optical faceplate, enabling a large field-of-view of up to 4 mm^3^ at a volume rate of 1 Hz. We demonstrate the power of this method with the first brain-wide recording of neuronal activity in an adult vertebrate.

## 1 Introduction

How brain-wide neural circuits operate is a fundamental open question in neuroscience. Answering it requires not only theoretical insight, but also adequate measurements. This has lead to a variety of optical imaging approaches, that, with help of genetically encoded calcium sensors, have enabled monitoring of neural activity of large populations of neurons. These techniques usually try to maximize their field of view, sampling rate and resolution in order to image a large number of neurons, located within a large region at high speed. As these parameters are interdependent all techniques are subject to specific trade-offs.

Optical neuroimaging experiments are often performed using two-photon point scanning microscopy [1]. Here, non-linear excitation is used to excite and collect fluorescence only in the vicinity of a laser focus, which is scanned through the specimen. Since the position of the laser focus as the origin of the fluorescence is known *a priori*, non-linear microscopy can deliver optically sectioned images. These techniques have been optimized to achieve large fields-of-view (FOVs) of several millimeter in diameter at diffraction limited resolution [2–5]. However, their speed is practically limited by the speed of the point-scanning and fundamentally by the fluorescence lifetime. Because excited fluorophores emit photons stochastically with a mean decay time on the order of nanoseconds, fluorescence from consecutive points could not be distinguished anymore, if measured at a higher rate.

Light sheet or selective plane illumination microscopy (LSM) circumvents these constraints by capturing entire planes of the specimen at once using a camera [6]. It employs one microscope objective to excite a thin sheet of fluorescence in the sample and an orthogonal imaging system to image this plane onto the camera. Because emission and excitation point spread functions (PSFs) are orthogonal, optical sectioning is maintained and volumetric imaging can be performed by consecutively scanning this plane through the specimen.

However, in many applications, optical access to the specimen from two orthogonal directions is not possible due to the specific mounting or occlusion by cartilage, bones, or other biological structures. Therefore light sheet imaging, so far, has been mainly used for neural imaging of transparent organisms like larval zebrafish [7–11].

Oblique plane microscopy (OPM), a variant of LSM, enables light sheet imaging even in the case of limited optical access. Here, oblique planes in the specimen are excited and imaged through the same objective [12–24]. However, the resulting image plane is oblique and therefore cannot be imaged onto a camera sensor using conventional approaches.

OPMs therefore employ a remote refocusing step in which an intermediate image of the oblique plane is created using a secondary microscope objective (Obj2). This intermediate image plane is brought to lie co-planar with the imaging plane of a tertiary detection objective (Obj3, Figure 1b). It can then be imaged onto a camera sensor.

**Fig. 1.**
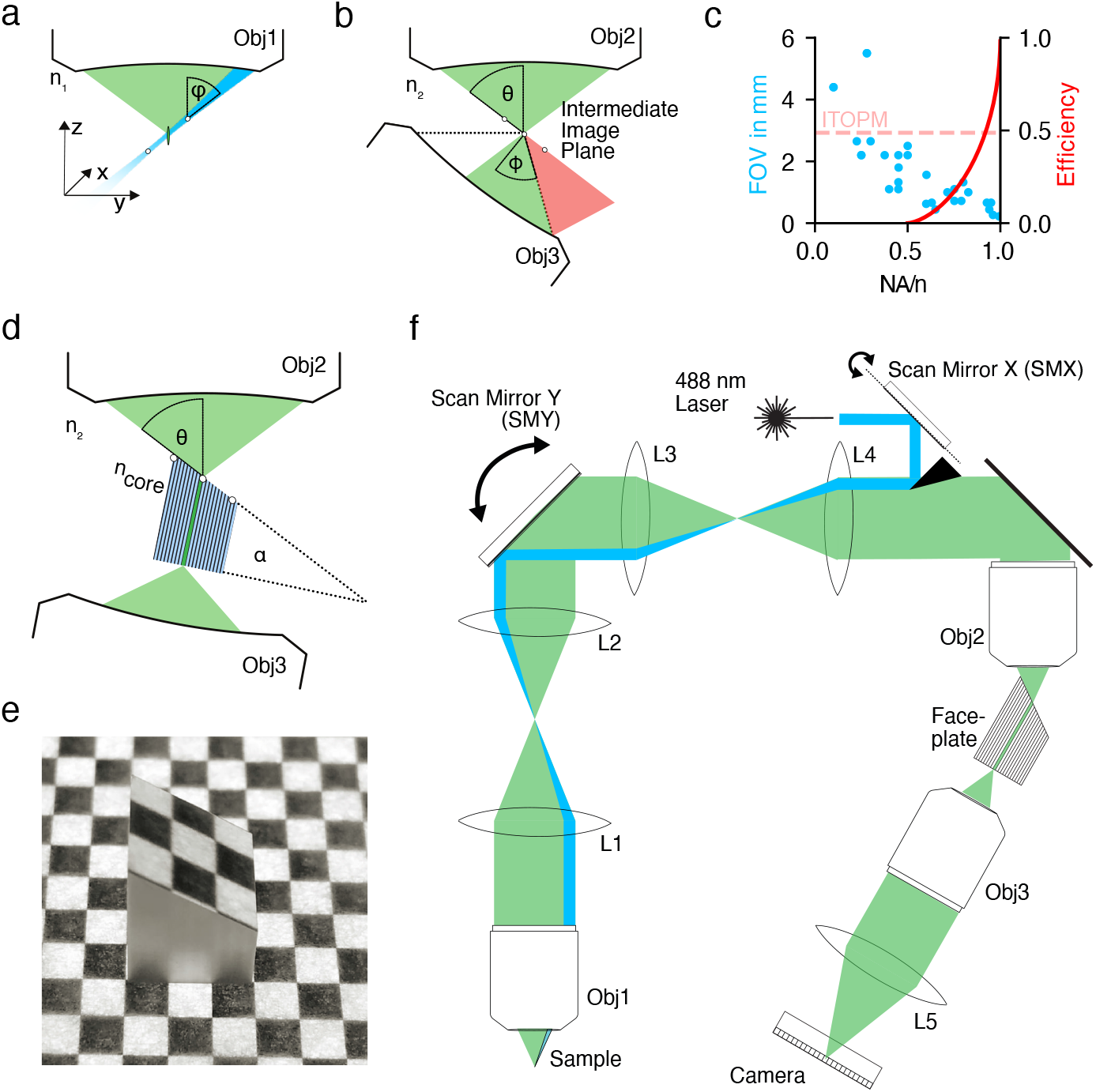
General idea and setup– a) Fluorescence is excited by a light sheet focused through the primary microscope objective (Obj1) at an angle φ. The image plane (white circles) is therefore oblique. Emitted fluorescence is then collected through the same lens. b) To image the oblique plane onto a camera, an intermediate image (white circles) is created by secondary objective (Obj2) at an angle *θ*. This plane is then brought to lie in the image plane of a tertiary microscope objective with an acceptance angle *ϕ*. This leads to loss of light for all *ϕ* < 90°. c) The efficiency of this re-imaging step is critically dependent on the NA of Obj2 and Obj3 (red curve). At the same time, the FOV scales inversely with the objective NA, shown in blue for some commonly used microscope objectives. d) Image transfer OPM employs a custom fiber optic faceplate (FP) with a core refractive index *n_core_*, that is cut at an angle *α*. *α* is chosen to minimize the coupling losses into the array of multi-mode fibers. The angled facet of FP is positioned at the intermediate image plane after Obj2. The intensity distribution at the intermediate image plane is then transmitted to the other end of the FP, where it is imaged by Obj3. e) A photograph of the FP placed on a printout of a checkerboard pattern. f) Complete setup consisting of the microscope objective lenses (Obj1-3), relay lenses (L1-L4, f=200mm), scan mirrors (SMX/SMY), the excitation laser, the fiber-optics faceplate and a high-speed camera

The magnification *M* between the specimen and the intermediate image plane, with refractive indices *n*_1_ and *n*_2_, is typically chosen to fulfill the condition of unit angular magnification 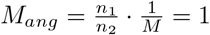 [25], resulting in *φ* = *θ* (Figure 1a, b). This ensures that first-order spherical aberrations are canceled [26] and even the points of the oblique plane that are outside of the native focal plane are imaged accurately.

The refocusing step, however, typically leads to a loss of light: Assuming an oblique plane at the limiting angle *ϕ* and identical objectives (*ϕ* = θ), some light will necessarily propagate outside of the half-angle of the acceptance cone of Obj3 *ϕ* if *ϕ* ≠ 90° (Figure 1b). This implies that this re-imaging solution cannot be used below NA/*n* = 0.5, as this loss becomes total. At the same time, due to constraints of optical design, the achievable FOV is inversely proportional to the NA of an objective (Figure 1c). This limits the achievable FOV of conventional OPM to about 1 mm (Figure 1c) [27, 28].

Multiple strategies have recently been proposed to mitigate this trade-off. A recent approach employs a microscope objective with an optically denser immersion medium at Obj3. This compresses the light cone and at NA=1 allows to capture close to 100% of the light [20, 22, 29]. Yet, Obj3 still needs to be a high NA objective to retain a substantial amount of light and therefore limits the FOV (to 300 (X) × 800 (Y) × 300 μm^3^ (Z) in [29]). Alternatively, the FOV of OPM can be de-coupled from the FOV of Obj3, by de-magnifying the larger FOV of a low-NA Obj1 onto the high NA intermediate image plane after Obj2 [21, 30], but the deviation from *M_ang_* = 1 rules out aberration-free imaging [26], limiting the axial imaging range or resolution. Without this de-magnificiation, refocusing below NA/*n* = 0.5 can be achieved by placing a diffraction grating at the intermediate image plane [31]. However, this reflective geometry is only compatible with low NAs (e.g. NA<0.3) that preclude cellular resolution.

Here, we present the first practical implementation of a novel OPM refocusing solution based on image transfer through an angled fibre optic faceplate, which works for low, intermediate, and high NAs (Figure 1 d,e). We demonstrate this by integrating our refocusing step into an OPM (Figure 1f), where it achieves refocusing with an efficiency of around 48% at an NA of 0.45. Using different primary objectives we achieve large FOVs of 2.4 × 2.1 × 1 mm^3^, while maintaining a resolution of at least 2.6 ± 0.7 × 3.1 ± 1.2 × 21.0 ± 5.7 μm^3^.

The uncoupling of NA from the efficiency of our refocusing step allows us to custom-tailor our design for adult brain-wide imaging in the small teleost fish *Danionella cerebrum* [32]. *Danionella* possess the smallest known vertebrate brain (2.5 × 1 × 0.8 mm^3^ [33, 34]) and remain transparent throughout adulthood [33–35]. Our microscope thus enables the first volumetric recordings of cellular activity across large parts of the adult vertebrate brain.

## 2 Results

### 2.1 A new approach for oblique plane re-imaging

Our design follows the basic principle of previous OPMs (Figure 1f) [12–24, 36]. At the heart of *image transfer oblique plane microscopy* lies a novel solution for the refocusing step of this oblique plane via a fiber optical faceplate (Figure 1c). This faceplate (FP, custom-made, material: 24AS, Schott AG, *n*_core_ = 1.81, *n*_cladding_ = 1.48, NA=1) is a rigid array of small multimode fibers with a diameter of 2.5 μm (Figure 1e). Therefore, it can transfer light intensity distributions from one plane to another without any additional optical system.

If the intermediate image plane of an OPM is brought to lie co-planar to the oblique surface of the FP, single fluorescence foci are coupled into the individual fibers of the FP. The opposite end of the FP can then be imaged onto a camera by the collection imaging system consisting of an objective (Obj3) and a tube lens (L5). Here the effective magnification is chosen to guarantee that each fibre was imaged at the Nyquist criterion.

The geometry of the FP needs to be optimized to ensure that most of the light is propagating within the coupling angle. During coupling, the light is refracted at the boundary between the imaging medium (air, *n*_2_ = 1) and the fiber core (*n_core_* = 1.81). We can cancel this refraction by selecting the apex angle *α* to be 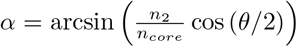 at 0°. This assumes that the mean incidence angle of the incoming cone is at half the acceptance angle *θ*. Here, the FP angle *α* = 35° was chosen to lie close to the optimal value 29°, to allow for some flexibility for alternative optical designs. We measured its light efficiency to be at 48% (See Methods).

### 2.2 Large FOV volumetric imaging

We can combine this re-imaging approach with different primary objectives (Obj1), which are close enough to the required unit angular magnification that ensures the cancellation of first-order spherical aberrations. Here we switched between an 10x/0.5 NA objective and a 16x/0.8 NA objective, with the goal to flexibly trade-off FOV at lower NA for increased axial resolution and collection efficiency at higher NA. This resulted in an effective angular magnifications of 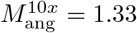 and 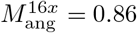.

To assess the performance of the microscope, we imaged a sample of fluorescent beads (*d* = 1μm) embedded in a polyacrylamide matrix with both primary microscope objectives. A Gaussian function was fitted through the line profiles along X,Y and Z of all detected beads.

As expected, the low NA variant achieved a higher FOV_10x_ = 2.4 × 2.1 × 1mm^3^, than the FOV_16x_ =2.1 × 1.7 × 0.8 mm^3^ (Figure 2a,f).

**Fig. 2.**
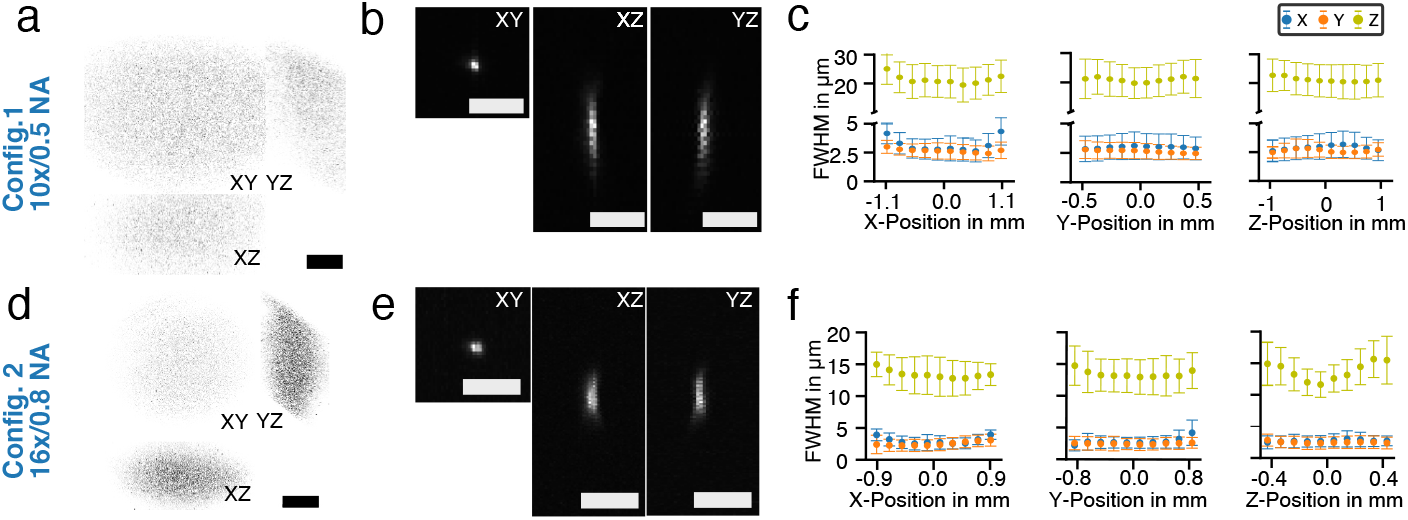
Resolution of the optical system – a) Orthoprojections (maximimum intensity projections, MIP) of a volume of fluorescent beads (d=1 μm, 2.4 × 2.1 × 1 mm^3^) imaged with the 10x configuration, b) orthographic MIPs of a single bead, c) mean and standard deviation of the measured full width at half maximum (FWHM, mean 2.7 ± 0.8 × 3.1 ± 1.1 × 21.1 ± 5.5 μm^3^ of beads across the FOV for all three axes show a relatively constant resolution. d) This procedure was repeated for the 16x configuration across a FOV of 2.1 × 1.7 × 0.8 mm^3^. e) The axial extent with the increased NA is smaller, f) which leads to an improved axial FWHM of 13.2 ± 2.8 μm, but increased axial variability. scalebars: a, d) 500 μm, b, e) 5 μm.

For the 10x configuration the full width at half the maximum (FWHM) was found to be 2.7± 0.8 × 3.1 ± 1.1 × 21.1 ± 5.5 μm^3^ (FWHM_10x_, n=17759, Figure 2 c). For the 16x microscope objective we measured 2.8 ± 1.0 × 2.4 ± 0.9 × 13.2 ± 2.8 μm^3^ (FWHM_16x_, n=17684, Figure 2)

Optical sectioning is important for the successful volumetric resolution of more densely labeled objects. We confirmed the optical sectioning capability of image transfer OPM by measuring the sum of the image counts in each axial plane of the image of a single bead. It was found to be 30 μm for the 10x and 11.3 μm for the 16x configuration (See Methods).

### 2.3 Brain-wide neuroimaging of *Danionella*

The flexible re-imaging of image transfer OPM allows us to optimize our microscope to specific neuroimaging applications and enables quasi-simultaneous recording over large volumes. Using the 16x configuration we demonstrate its practicability by imaging the brain of *Danionella*, the smallest known vertebrate brain [33] of typically 2.5 mm × 1 mm × 0.8 mm. We therefore could accommodate almost the entire brain of adult *Danionella* with pan-neuronal nuclear expression of GCaMP6s (huc:h2b-gcamp6s x tyr -/) in the FOV of 2.1 × 1 × 0.8 mm (Figure 3a)

**Fig. 3.**
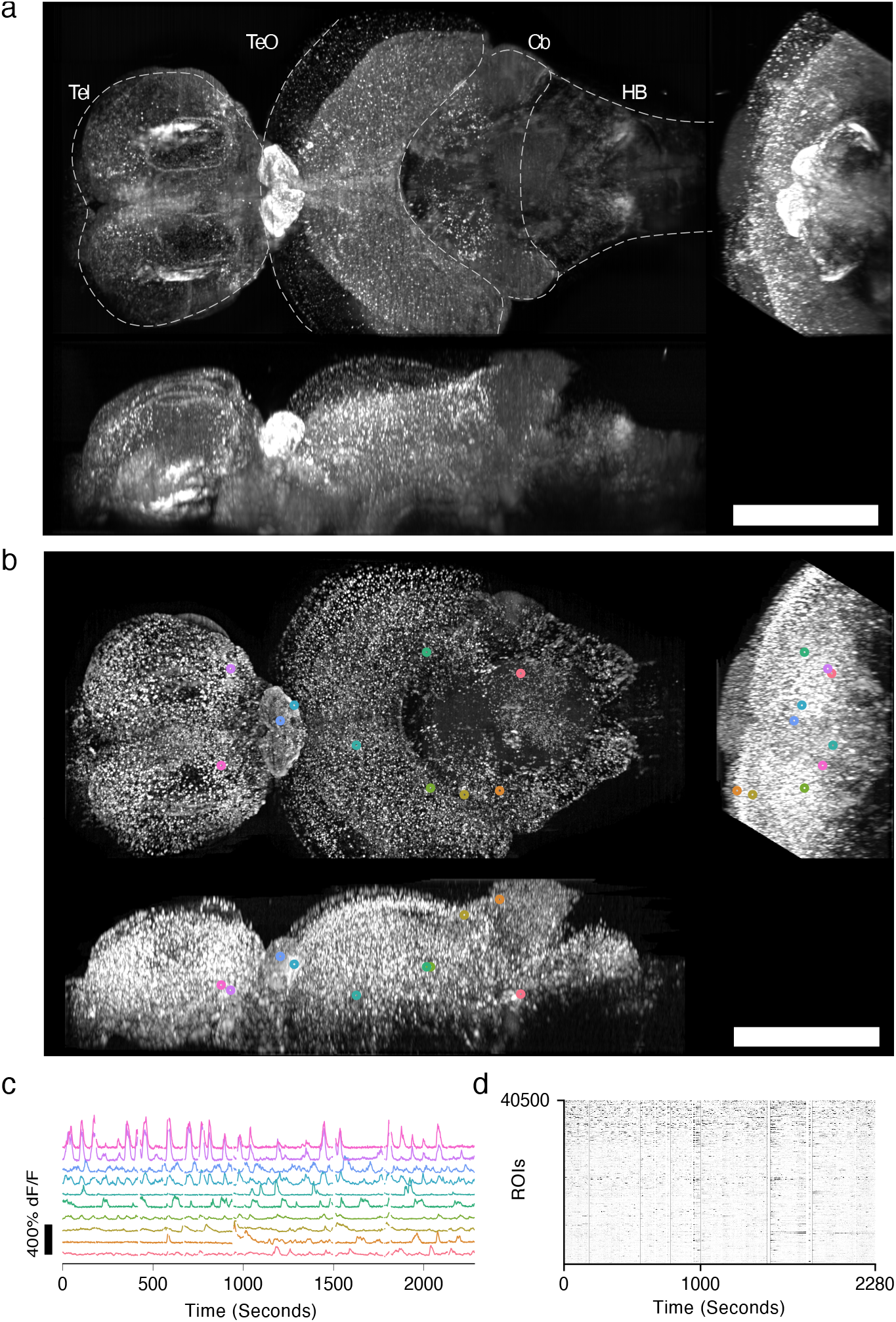
Brain-wide imaging of neural activity in adult *Danionella* - a) Orthoprojections (maximum intensity) of the time series mean of an adult *Danionella c*. (huc:h2b-gcamp6s x tyr -/-). The recording lasted 38 min at a volume rate of 1Hz. The FOV encompasses many major brain regions, such as the telencephalon (Tel), optic tectum (TeO), cerebellum (Cb) and hindbrain (HB). b) Local temporal correlation map serving as the segmentation map of the brain. c) Fluorescence dynamics from ten neurons at different locations marked in a) throughout the entire brain. d) The nuclear expression allowed us to segment ≈ 40k putative neurons by local maxima detection and extract their time dependent fluorescence (white bands: excluded frames due to motion artefacts). Scale bar: 500 μm

Recording 318 Y-planes (1000×3000 pixel) per second at a spacing of 3 μm allowed us to image the brain at 1 Hz. These designs allowed us to quasi-simultaneously record neural activity from 44k ROIs throughout distant brain regions, such as the brainstem, cerebellum, telencephalon, and thalamus in an adult vertebrate organism (Figure 3b,c,d).

## 3 Discussion

Image transfer OPM makes use of a novel refocusing solution based on an angled fiber-optic faceplate to achieve multi-millimetre FOVs at cellular resolution. In contrast to conventional OPM [12], whose efficiency approaches 0 as the NA/n decreases towards 0.5, our approach allows us to refocus the oblique plane at all NAs. We therefore are able to choose primary objectives with intermediate NA (10x/NA 0.5, 16x/NA 0.8) and to achieve imaging volumes up to 4 mm^3^ (FOV_10x_ = 2.4 × 2.0 × 1.0 mm^3^, FOV_6x_ = 2.1 × 1.7 × 0.8 mm^3^), larger than in previous high-NA OPMs [15, 17, 22, 37]. At the same time, image transfer OPM maintains a better resolution than previous large FOV OPMs that employed low NA objectives [31, 38].

The efficiency of our re-imaging system is 48%, determined by coupling efficiency, Fresnel reflection losses and fill-factor. This is equivalent to a conventional OPM [12] refocusing efficiency at about 0.83 NA/n (Figure 1). It could be increased with the help of anti-reflective coating.

Being a one-photon camera-based technique, image quality is reduced by scattering effects as we image deeper. To improve image quality a camera-based line confocal detection scheme [39, 40] or a structured illumination approach could be integrated. As our laser sweeps across the plane to create the excitation sheet, it could be modulated, and several differently illuminated images could be computationally combined to yield a higher resolution [41]. The resolution could be further improved by realizing our system as a dual-view microscope [42] similar to [29] or with an additional imaging and illumination arm for different perspectives.

In our microscope, we magnify the surface onto the camera in order to avoid Moiré effects between camera pixel (2.5 μm) and fiber grid (2.5 μm). Alternatively, the FP could be directly attached to the sensor surface if the pixel grids were matched. Therefore, our re-imaging unit could be thought of as a first approximation to an image sensor for oblique planes rather than a tertiary imaging system. With further miniaturization of pixels of scientific cameras and the development of nanophotonic structures for efficient detection of oblique light, this direct sensing could become the default.

The impact of our development is hopefully twofold. First, we demonstrate how the re-imaging efficiency of an OPM microscope can be decoupled from the numerical aperture. This permits the construction of OPMs at virtually any NA, enabling highspeed volumetric imaging at large FOVs.

Second, our microscope has unique capabilities of immediate use in neuroscience. In particular, the combination of image transfer OPM with the advantages of the transparent model organism *Danionella* enables the first volumetric recording of brain-wide neural population activity in any adult vertebrate. This expanded capability of optical neural population imaging will lead to new insights into the properties and function of the nervous system at a systemic level.

## 4 Online Methods

### 4.1 Optical Setup

As typical for an OPM, our design essentially consists of three imaging systems. As the primary microscope objective (Obj1) facing the specimen, we employ either a 10x (*f* = 10 mm, NA = 0.5, water-immersion, CFI Plan Apochromat 10XC Glyc) or a 16x (*f* = 12.5 mm, NA = 0.8, water-immersion, 16X CFI LWD Plan Fluorite, Nikon) objective lens. The excitation laser (06-MLD, 488 nm, Cobolt) hits a scanning mirror (SMX, 6 mm, 8315K, Cambridge Technology) and is subsequently reflected into the microscope via a pick-off mirror (PM). The center of SMX is imaged onto the back-focal aperture (BFP) of Obj1 (Obj1) via two 4-f systems (L1-L2, L3-L4, all TTL200MP, Thorlabs Inc., f=200 mm). SMX is positioned off-axis. A fast-scanning SMX, therefore, creates an oblique light sheet within the specimen, exciting fluorescence. The laser beam forming the light sheet had a waist of *ω*_10*x*_ = 7.9 μm and *ω*_16*x*_ = 7.3 μm resulting in a calculated Rayleigh range of 400 μm and 350 μm respectively. The angle of the oblique plane in the specimen was *φ*_10*x*_ = 21° and *φ*_16*x*_ = 33°. This oblique plane is then imaged onto the intermediate imaging plane of Obj2 (CFI Plan Apo Lambda 10X, *f* = 10 mm, NA = 0.45) at an angle of θ = 27°.

If the intermediate image plane of an OPM is brought to lie co-planar to the oblique surface of the faceplate (FP), single fluorescence foci are coupled into the individual fibers of the FP. The opposite end of the FP can then be imaged onto a camera (CB262CG-GP-X8G3, Ximea, pixel pitch = 2.5 μm, 5120×5120 pixel) by the collection imaging system consisting of an objective (Obj3, CFI Plan Apo Lambda 10X, *f* = 20mm, NA = 0.45) and a tube lens (L5, XLFLUOR 4X, *f* = 45mm, NA = 0.28, Olympus). The effective magnification of 2.5 guarantees that each fibre was imaged at the Nyquist criterion.

Volumetric imaging is enabled by a second galvanometric mirror (SMY, 25 mm diameter, 6240H, Cambridge Technology), which is conjugated to the BFP of the primary objective and allows to scan the light sheet throughout the specimen and to de-scan the emission light onto the static intermediate image plane. The synchronization of scanning and camera image acquisition are controlled via custom software (Python) and a data acquisition card (NI USB-6363, National Instruments).

All camera images are recorded with an exposure time of 3 ms with an additional read-out time of 0.145 ms. Our camera has a 10-bit range at the read-out layer. The acquired data is mapped onto a 8-bit range before being transferred to the host computer using a custom linear LUT that fixed the gain to around 5*e*^-^/count. This allows us to reach higher frame rates.

### 4.2 Characterisation of image transfer efficiency

At the heart of ITOPM lies a re-imaging system that employs a fibre-optical faceplate (FP) to re-focus the oblique image plane. This FP consist of multimode fibres (NA 1.0, diameter 2.5 μm, core-cladding ratio 0.7). In order to characterise the efficiency of this re-imaging step we used a green laser (532 nm, MGL-DS-532, CNI). The laser was coupled into a single-mode fiber (P3-405B-FC, Thorlabs). The output of the fiber was then collimated using a microscope objective (2X, 0.1 NA, 56.3 mm WD, Thorlabs), with the same 20 mm back focal aperture as objective Obj2. This collimated output was then coupled into the system at the back focal aperture of Obj1. The intensity of the resulting laser focus at the intermediate imaging plane was measured using our tertiary imaging system without the FP. We then inserted the FP and measured the intensity of the light as it would be imaged with ITOPM. The tertiary imaging system was positioned such that the laser was spread out over multiple fibre cores at the front of the FP to account for the fill fraction.

### 4.3 Characterisation of resolution

Fluorescent beads (diameter 1 μm) were dispersed in a poly-acrylamide gel between a glass slide and a coverslip, separated by a silicon spacer. A stack of the whole accessible image volume was taken. The stack was then preprocessed as described above, but not deconvolved. The stack was thresholded at the 99.99th percentile and all connected components were segmented out within a ROI of 33 × 33 × 303 μm^3^ (X × Y × Z) For the quantification of the lateral resolution each ROI was maximum intensity projected along z. The lateral resolution in x and y were determined as the full width at half maximum (FWHM) of the Gaussian fit through of the line plot along the maximum of this projection. The axial sectioning capability was determined as the FWHM of the Gaussian fit of the sum of all pixel values along the XY planes of each bead volume.

### 4.4 Postprocessing

After the data is acquired several postprocessing steps are executed. For every plane the camera background is substracted. Subsequently grid artefacts, that stem from the surface structure of the face plate are corrected. A reference grid image is obtained by imaging a homogeneously fluorescent object. Each camera frame is then corrected by dividing the frame by the correction pattern.

Each volume

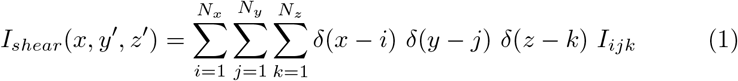

is natively recorded in a sheared coordinate system by our microscope and needs to be unsheared for convenient analysis. During this process we simultaneously execute two other steps: 1) We axially bin our volumes along *z* = *z*’ since the effective pixel pitch of the camera (dz’=0.73 μm) is smaller than the axial resolution. 2) We upsample the volume along the *y*-direction to 1 μm from the native sheared Y’-direction, which is originally sampled at 3.5 μm. This is necessary and beneficial since the projection of the microscope PSF onto the sheared Y’, a mixture of axial and lateral resolution, is larger than onto the Y. During unshearing we can therefore re-cover intermediate Y planes through interpolation.

This is done by computing the unsheared upsampled and axially binned volume at a new coordinate grid with voxel size 0.75 × 1.0 × 4.5 μm^3^ (XYZ) by using the following interpolation kernel

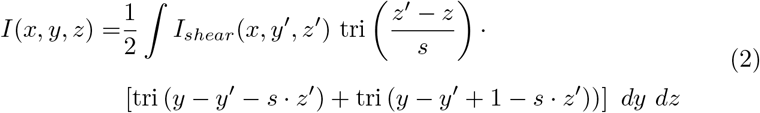

Here, tri(*x*) = *max*(0, 1 – ||*x*||) and *s* is the slope of the oblique plane.

Lastly, in the case of neuro-imaging data, we deconvolve the data using 10 iterations of Richardson-Lucy deconvolution. The kernel we used is empirically estimated from an average of previously imaged fluorescent beads.

### 4.5 Neuroimaging of *Danionella cerebrum* (Dc)

Dc specimens were kept in commercial zebrafish aquaria (Tecniplast) with the following water parameters: pH 7.3, conductivity 350 μS/cm, temperature 27 ^°^C. Adult Dc, expressing an histone-tagged GCaMP6s pan-neuronally (huc:h2b-gcamp6s x tyr -/-) were embedded in a pre-formed agarose mold, which allowed the gill covers to move freely, and immobilized with 2% low-melting point agarose. A mouthpiece, made from a glass pipette was inserted into their mouth and the fish were then perfused with fresh aquarium water via a peristaltic pump. They were allowed to recover from anesthesia for 15 minutes prior to experiment onset. After experiment onset the intensity of the excitation beam was gradually increased over 2 minutes up to a final power of ≈5.3 mW after the imaging objective to allow for slow habituation. The beam was scanned through one plane in 2 ms and coincided with the 2 ms sensor exposures. Recording and read-out of one plane took 3.1 ms. We could therefore image 332 planes spanning 827.5 μm at 1 Hz volume rate.

### 4.6 Image registration

In order to analyze timelapse recordings of whole-brain imaging datasets all volumes had to be motion corrected. The dataset processed for this manuscript consisted of 2400 volumes with a size of 3024 x 960 px x 144 px (XYZ).

One volume from the recording was selected as the template, on to which all other volumes were registered.

Before estimation all volumes were band-pass filtered with a difference of Gaussian filter (σ_0_ = 2 px, σ_1_ = 5 px) to enhance high frequency features and exclude residual grid artefacts introduced by the face plate.

The registration consisted of the estimation of an affine and a non-rigid transformation from estimates of locally rigid displacements.

Practically, each volume was chunked into non-overlapping blocks and the three dimensional rigid displacement for each of block was then determined via cross-correlation.

In a first iteration, a global affine transformation was fitted onto the obtained coarse displacement field with a blocksize [604 px, 192 px, 28 px] (XYZ). After correction of the coarse affine warp, the volume was again divided in to smaller blocks of [32 px, 32 px, 16 px] (XYZ) to estimate a better resolved displacement field. From this field, a full non-rigid displacement field was then obtained via interpolation. Finally the compound transformation consisting of affine and non-rigid transformation was applied onto the original volume using linear interpolation.

### 4.7 Source extraction

To extract neural activity from the data a simple local maxima detection was used. The local correlation of each voxel with its neighbourhood (26-connectivity) was calculated. Before doing so, frames within 10 seconds before and after spontaneous animal motion were dropped to exclude motion artefacts. Local maxima within spherical surroundings (d=10 μm) that corresponded to the expected size of the labeled cell nuclei were detected. Afterwards the number of local maxima was plotted against the threshold, given as the intensity percentile. This curve had an inflection point, which was used as the final threshold. Additionally, the volume was masked to exclude eyes and edge artefacts. Each volume was then convolved with a sphere of diameter 10μm, that corresponded to the expected size of the labeled cell nuclei. The resulting intensity trace at each local maximum was then extracted as the fluorescence trace of this point. To compute the dF/F0 trace a baseline fluorescence F0(t) was computed by median filtering F(t) (windows size = 120 s), minimum filtering (windows size = 120 s), and then smoothing (Gaussian smoothing, windows size = 120 s) F(t). dF/F0(t) was then computed as dF/F0(t)=(F(t)-F0(t))/F0(t).

## Supplementary information

## Acknowledgments

The authors thank Caroline Berlage, Thomas Chaigne, Verity Cook, and Fabian Voigt for critically reading the manuscript.

Data analysis was performed on the Berlin Institute of Health high-performance compute cluster.

## Declarations

### Ethics approval

All animal experiments conformed to Berlin state, German federal and European Union animal welfare regulations and were approved by the LAGeSo, the Berlin admission authority for animal experiments.

### Availability of data and materials

Data used for analysis will be made available at a g-node repository.

### Code availability

All analysis code will be made available at www.github.com/danionella/itopm

